# From Diversity to Discovery: Genome-Wide Insights into the Genetic Landscape of Tropical Maize DH Lines

**DOI:** 10.64898/2026.02.03.703488

**Authors:** Azalech Oli, Solomon Benor, Gizachew Haile, Berhanu Tadesse, Yoseph Beyene, Manigben Kulai Amudu, Manje Gowda

## Abstract

Understanding the extent and structure of genetic diversity within breeding populations is essential for sustaining long-term genetic gain in maize improvement programs. In this study, a panel of 2,555 maize doubled haploid (DH) lines representing diverse genetic backgrounds was genotyped using 3305 high-quality single nucleotide polymorphism (SNP) markers to assess genome-wide diversity, population structure, and relatedness. The SNPs were distributed across all ten chromosomes, with varying marker densities among genomic regions. Diversity indices revealed moderate polymorphism, with mean gene diversity (0.38) and polymorphic information content (0.30), while the minor allele frequency ranged from 0.04 to 0.50. The low observed heterozygosity (0.04) and high fixation index (0.89) confirmed the expected homozygosity of DH lines. Population structure analysis using sparse non-negative matrix factorization (sNMF) and principal coordinate analysis (PCoA) consistently identified two major genetic clusters corresponding to the established heterotic groups used in CIMMYT’s tropical maize breeding pipelines. The Analysis of Molecular Variance (AMOVA) indicated that 36% of genetic variation occurred among populations, 58% among individuals within populations, and 6% within individuals (*P* = 0.001), confirming significant population differentiation and high within-group diversity. These results demonstrate that the DH panel represents a genetically diverse and well-structured population with limited relatedness among lines. The distinct clustering by heterotic group, coupled with substantial within-group variation, provides a strong foundation for genome-wide association studies, genomic selection, and allele mining for complex adaptive traits. The panel’s diversity and structure make it an invaluable genomic resource for dissecting trait architecture and accelerating genetic gain in tropical maize breeding programs targeting sub-Saharan Africa and similar environments.

## Introduction

Maize (*Zea mays* L.), a member of the Poaceae family, is domesticated about 9,000 years ago in southern Mexico and Mesoamerica [24]. Today, it has grown at more than 200 million hectares worldwide and ranks as the third most widely cultivated cereal after wheat and rice [12]. Because of its high yield potential and extraordinary versatility, maize is widely regarded as the “queen of cereals” [41]. In sub-Saharan Africa (SSA), maize is the most important staple crop, contributing more than 30% of daily caloric intake [41]. It is cultivated across highly heterogeneous agroecological zones, ranging from the humid lowlands of West Africa to the drought-prone mid-altitudes of Eastern and Southern Africa. Despite its wide adaptation, maize productivity in SSA remains low, averaging less than 2 t ha^−1^ compared with global averages of more than 5 t ha^−1^ [14]. With a rapidly growing population and changing consumption patterns, the demand for maize in SSA is projected to increase substantially, with current estimates indicating a production-consumption gap of about 45% [12]. This gap highlights the urgency of accelerating genetic improvement and enhancing productivity per unit area.

Several constraints limit maize productivity in SSA. Biotic stresses include parasitic weeds such as *Striga hermonthica*, insect pests such as fall armyworm (*Spodoptera frugiperda*), and diseases including maize lethal necrosis, turcicum leaf blight, and gray leaf spot [33]. Abiotic stresses, especially drought and low soil fertility, further reduce productivity, while climate variability adds unpredictability to farming systems. These constraints are exacerbated by the predominance of smallholder farming systems with limited access to inputs such as fertilizers and irrigation. Meeting these challenges requires maize varieties with enhanced resilience, stability, and yield potential under stress-prone environments [4].

The success of hybrid maize breeding relied heavily on heterosis or hybrid vigor. Exploiting heterosis requires a systematic understanding of genetic diversity, population structure, and heterotic group formation [25, 17]. Doubled haploid (DH) technology has transformed maize breeding by enabling the development of fully homozygous inbred lines within just two generations, compared with five to eight generations required by conventional selfing [35, 5]. CIMMYT and national research partners have widely adopted DH technology in SSA, allowing rapid generation of large numbers of inbred lines for hybrid testing. The ability to produce DH lines accelerates the breeding cycle, enhances selection accuracy, and provides a rich reservoir of parental lines for genetic diversity studies and hybrid development.

The assessment of genetic diversity and population structure is central to harnessing the potential of DH lines and other germplasm resources. Genetic diversity enables breeders to identify parental lines carrying complementary alleles for yield, stress resilience, and quality traits [43,39]. Knowledge of population structure is critical to define heterotic groups, design efficient crossing strategies, and maximize heterosis in hybrid breeding [21]. Furthermore, genetic diversity studies help maintain a broad genetic base, reducing the risks of genetic vulnerability to pests, diseases, and climate variability [46].

Molecular markers greatly improved the resolution of genetic diversity and population structure analyses in maize [27]. Early studies employed markers such as restriction fragment length polymorphisms (RFLPs), amplified fragment length polymorphisms (AFLPs), and simple sequence repeats (SSRs) [42, 40]. These markers provided valuable insights into genetic variability in tropical and temperate maize germplasm but were limited by low throughput and incomplete genome coverage. The advent of high-throughput single nucleotide polymorphism (SNP) genotyping has enabled genome-wide characterization of maize diversity with unprecedented precision [11,47]. SNP markers are abundant, co-dominant, locus-specific, and cost-effective, making them the marker of choice for large-scale diversity studies. SNP-based studies have been instrumental in classifying tropical maize germplasm into heterotic groups and guiding parental selection. [5] used nearly 100,000 SNPs to assign 417 tropical maize DH lines into heterotic groups, while [16] applied SNP data to characterize 310 DH lines for oil content improvement. Such studies demonstrate the power of SNPs in uncovering population structure and identifying optimal parental combinations for hybrid development.

For SSA, understanding genetic diversity and population structure has both theoretical and applied significance. Theoretical insights into the partitioning of genetic variance within and among populations strengthen knowledge of tropical maize evolutionary history and adaptation. Practically, these analyses support hybrid breeding by informing heterotic group assignments and ensuring efficient use of germplasm collections maintained by CIMMYT and IITA. Genetic characterization also helps safeguard against genetic erosion by identifying lines that contribute unique alleles for stress tolerance, nutritional quality, and other farmer-preferred traits. Such efforts are crucial for developing resilient, high-yielding hybrids tailored to SSA’s low-input, stress-prone farming systems.

Modern statistical tools further enhance the utility of diversity studies by facilitating robust heterotic group formation. Classical methods such as diallel and testcross approaches have long been used to identify heterotic patterns [38,18]. However, these approaches are resource-intensive and limited in their ability to handle large germplasm sets. The integration of advanced molecular marker data with statistical clustering, principal component analysis, and model-based population structure analysis provides an efficient, scalable alternative for assigning large numbers of inbred or DH lines to heterotic groups [36].

Given the critical role of maize in SSA’s food systems, the urgent need to close the region’s production-consumption gap, and the transformative potential of DH and SNP technologies, comprehensive assessments of genetic diversity and population structure are essential. The present study leverages a unique panel of 2,555 DH lines derived from 87 biparental populations developed within CIMMYT’s tropical maize breeding program. Using 3305 SNPs generated through DART mid-density platform, we aimed to (i) quantify the extent of genetic diversity within and among DH lines, (ii) characterize population structure and heterotic patterns, and (iii) assign lines to heterotic groups relevant for hybrid breeding in SSA.

## Materials and Methods

### Plant material

A total of 2,555 DH lines derived from 87 bi-parental population (Table 1, Supplementary Table S1) were used in this study. The base germplasm from which these 2555 DH lines were generated was developed by the International Maize and Wheat Improvement Center (CIMMYT) for mid-altitude environments (referred to as CIMMYT base germplasm). These DH lines were pre-selected for their high yield, resistance to maize lethal necrosis, maize streak virus diseases and tolerance to drought stress. These DH lines were part of stage I (early-stage yield testing in breeding pipeline) lines used in a subset from 2017 to 2021 in the early to medium maturity breeding pipeline of eastern Africa. All these DH lines were generated at the CIMMYT-Kenya DH facility following the procedure described by [34]. The 87 biparental populations were derived from CIMMYT eastern African mid-altitude germplasm with different genetic backgrounds.

### DNA extraction and sequencing

Seeds of the DH lines were first planted in a greenhouse. Three weeks after planting, four leaf discs were punched from the leaves, placed in a deep-well plate leaf samples were collected and stored at −80 °C for 72 hours. The frozen samples were then dried using a Labconco Freezone 2.5L lyophilizer (Marshall Scientific, USA) and sent to Intertek Laboratory in Australia for mid-density DArTseq genotyping. The genotyping process followed the method described [20]In brief, DNA was first extracted from the leaf samples, purified, and cut with restriction enzymes at specific target sites. Short DNA adapters with unique sequences were then attached to the cut fragments. Next, a proprietary amplification method was used to reduce genome complexity and enrich for regions containing the selected SNP markers. PCR was then carried out to produce millions of copies of these fragments, which were used for SNP analysis. The amplified DNA was hybridized with probes, imaged, and analyzed on the DArT platform to identify the presence or absence of SNPs in each sample. Finally, data quality control and genotyping calls were performed following [20].

### Marker quality control

The DArT platform generated 3,305 SNP markers through its analytical pipeline. To ensure reliability, marker quality control was done in TASSEL version 5 (Bradbury, et al. (2007) using several criteria: minor allele frequency (MAF ≥ 0.05), call rate (≥ 90%), missing data rate (≤ 0.25), genotype quality (GQ = 20), bi-allelic SNPs only (no indels), and two alleles per locus. After filtering, 2,257 high-quality SNPs remained and were used for downstream analyses.

### Genetic diversity and population structure analyses

The final HapMap with 2,257 high-quality SNPs was converted to variant call format (VCF). Using PLINK 2.0 [6], we estimated genetic diversity indices, including expected and observed heterozygosity (EH, OH), minor allele frequency (MAF), and polymorphism information content (PIC). Additional diversity measures such as Shannon index, inverse Simpson index, and Alpha index [7] were also calculated to quantify allele richness and evenness. The HapMap file was further converted to MAF format, from which a gene content file was generated in R [37].

The adegenet, ape and scales packages within the DartR platform were used for Discriminant Analysis of Principal Components (DAPC), Principal Coordinate Analysis (PCoA) and Hierarchical Clustering of the DH lines. The clustering results were compared with the clusters generated using the Bayesian Information Criterion (BIC). The filtered and SNP data were analyzed with R software version 4.4.1. Pairwise population differentiation matrices among genotypes were calculated using GenAlEx [31]. The analysis of molecular variance (AMOVA) was conducted by GenAlEx, version 6.51b2 software. A clean dataset of 2,555 DH lines was analyzed to estimate kinship values based on identity by state (IBS) using R software version 4.4.1. Also, to assess group differences and the heterotic alignment between the DH line based on structure.

The genetic structure of the maize DH panel was analyzed using the **LEA v3.4.0** package in R. We used one statistical approach—sparse non-negative matrix factorization (sNMF) employed to estimate population ancestry and visualize genetic relationships among lines. The *snmf* function was executed for K values ranging from 1 to 30, with 10 replicates per K and a regularization parameter of α = 100, to identify the optimal number of ancestral populations. The LEA algorithms are particularly well suited for estimating ancestry coefficients in inbred or selfing lineages, making them advantageous over programs such as ADMIXTURE for datasets like the current DH panel. Complementary to this, PCoA was performed using the adegenet R package to further explore the genetic structure and clustering patterns within the population. Analysis of molecular variance (AMOVA) and genetic differentiation (F_ST_) was done using GenAlEx 6.5 [20] to partition the variation between and within populations.

## Results

The Supplementary Table S1 provides pedigree information for the DH maize lines used in the study. These lines are classified into heterotic groups A and B, reflecting breeding origins and relationships within the material tested. This foundational data supports subsequent diversity analyses by contextualizing genetic variation with respect to known backgrounds. In this study, doubled haploid lines of maize were genotyped using 3305 informative SNPs with a call rate of greater than 0.2. Out of 3,305 SNP markers, monomorphic markers and those with minor allele frequency of less than 0.05 were removed, resulting in 2257 high-quality SNP markers being used for further analysis. A total of 2,257 high-quality SNP markers were successfully mapped across the ten maize chromosomes in 2,555 DH lines. The SNPs were unevenly distributed along the chromosomes, with some regions showing dense marker coverage (7–8 SNPs per segment) and others more sparsely represented (1–3 SNPs) (Figure 1). This genome-wide distribution ensured adequate representation of the maize genome for downstream diversity and structure analyses. The markers exhibited a broad range of allele frequencies with a mean minor allele frequency (MAF) of approximately 0.30. This distribution indicates a population with moderate allelic variation, including a balance of common and rare alleles across the genome.

**Figure 1.**
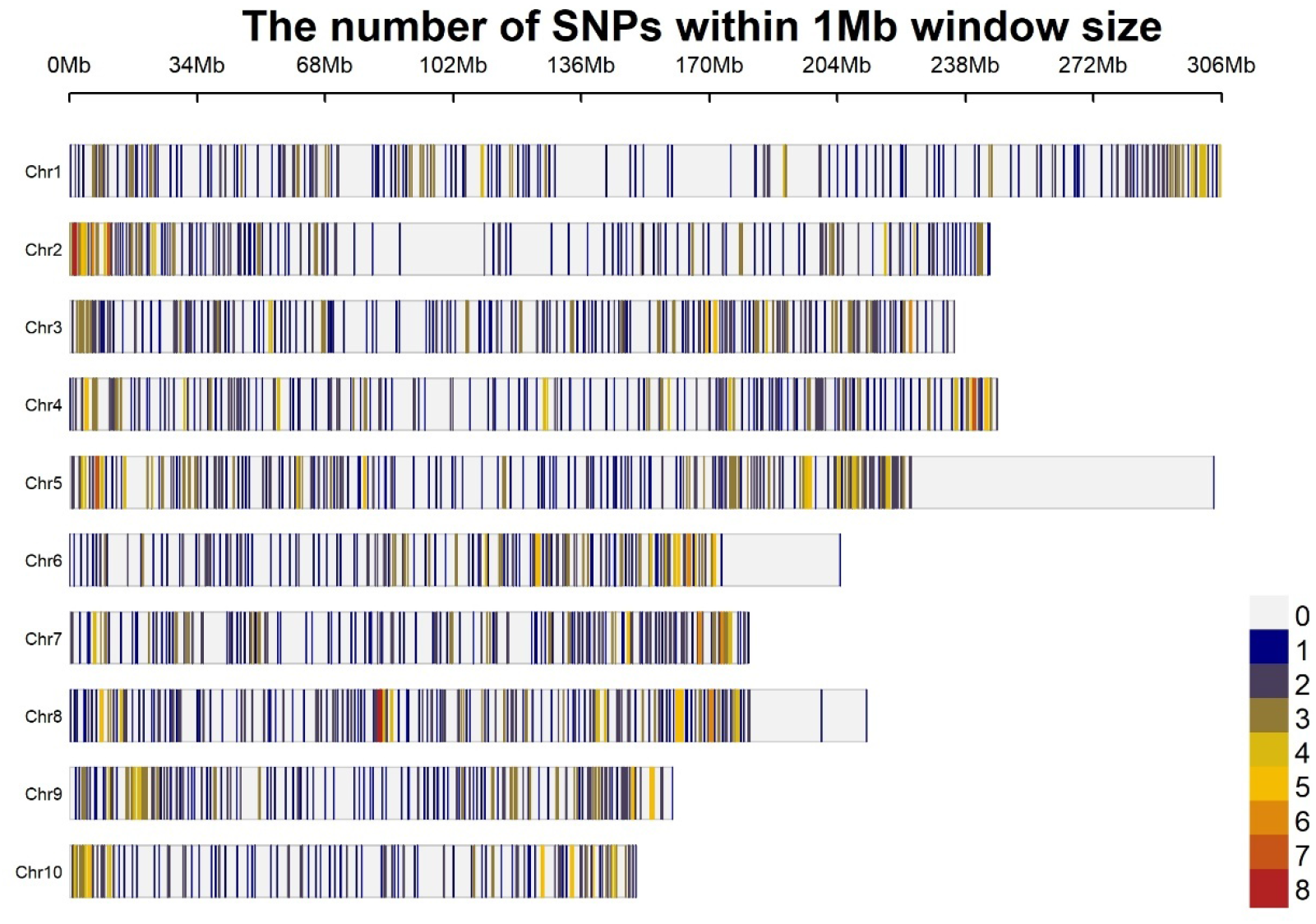
Distribution of 2257 SNPs across the 10 chromosomes of the maize genome. The number in the key indicated the quantity of SNPs corresponding to the color pattern depicted in the plot. Grey; no markers, dark to light blue; 1–3 SNPs, light to dark yellow and orange; 4–6 SNPs, light and deep red; 7 and 8 SNPs, respectively.

The expected heterozygosity (He), which represents the theoretical genetic variation assuming Hardy-Weinberg equilibrium, averaged 0.38 (Figure 2), signaling a moderately diverse germplasm. In contrast, the observed heterozygosity (Ho) was substantially lower (∼0.04), which aligns with the use of inbred or doubled haploid lines where homozygosity is intentionally fixed through breeding processes. This discrepancy underscores the impact of breeding and selection on maize genetic makeup, fostering genetic uniformity within lines while retaining allele variation at the population scale.

**Figure 2.**
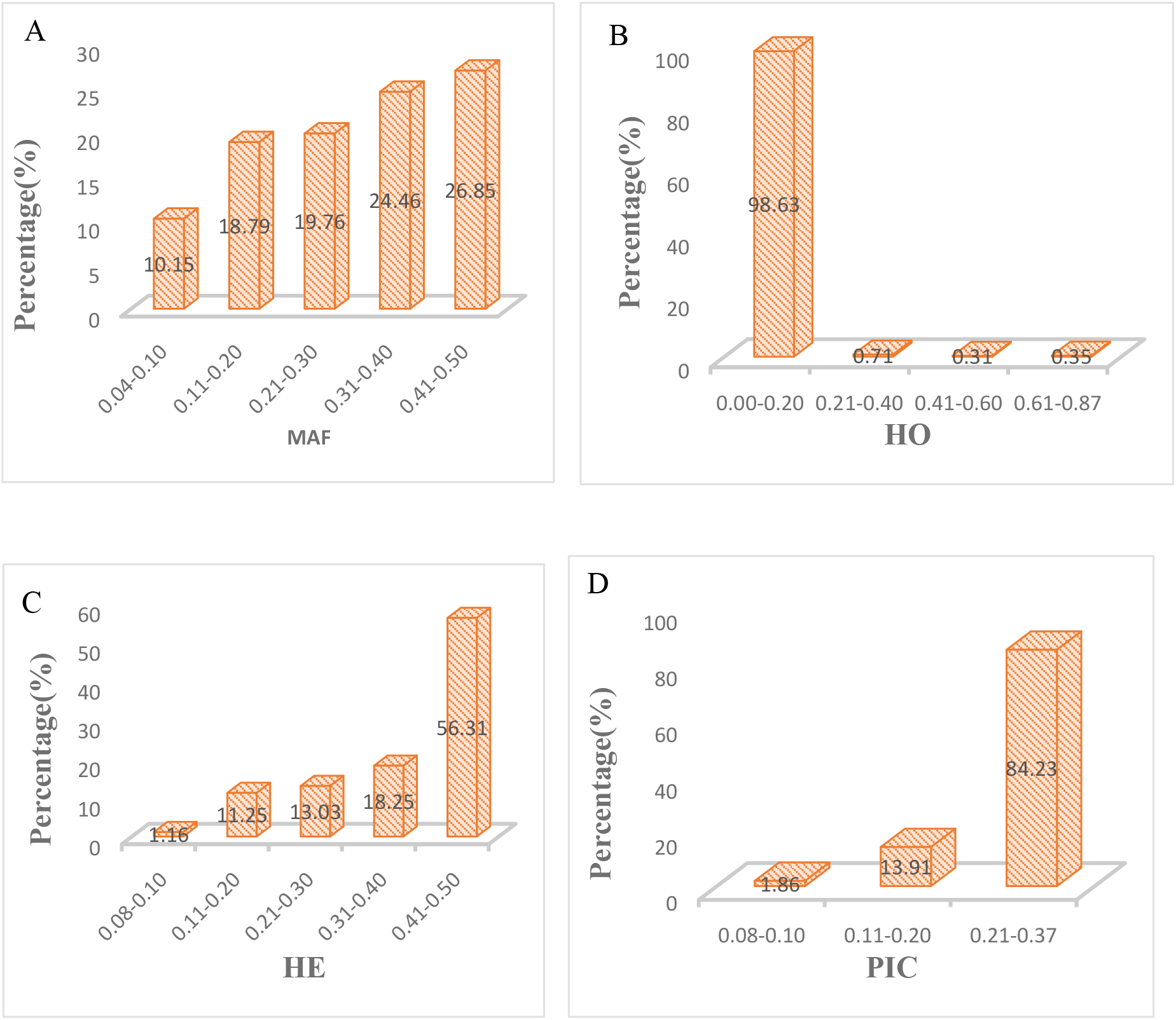
Summary of 2257 SNP markers with 2555 maize DH lines genotyping of: (A) Minor Allele Frequency (MAF), (B) Observed Heterozygosity (Ho) (C) Expected Heterozygosity and Polymorphic Information Content (PIC).

Polymorphism Information Content (PIC), a measure assessing marker informativeness for genetic diversity and linkage studies, averaged 0.30 (Figure 2). This metric validates the selected SNP marker set’s suitability for capturing population diversity and structuring analyses. Complementing these diversity measures, the inbreeding coefficient averaged 0.89, confirming the high homozygosity characteristic of the panel. This high inbreeding level is expected in doubled haploid and inbred maize populations and reflects the intensity of artificial selection during line development.

The effective population size (Ne) was estimated at around 1432, indicating a sizable genetic base that facilitates maintaining diverse alleles within breeding pools. Genetic variance components revealed substantial additive genetic variance (∼863), implying a strong potential for selection response, along with notable dominance variance (∼362), important for heterosis breeding considerations. Together, these analysis results paint a clear picture of the maize diversity panel as one with considerable allele diversity but highly homozygous individual lines. This supports the use of the panel for genetic mapping, association studies, and breeding applications aimed at improving important agronomic traits while maintaining genetic diversity essential for long-term crop improvement. The detailed pedigree information, combined with robust population genetic analyses, enhances the understanding of maize genetic architecture in the studied material and underpins the molecular breeding approaches for resistance and yield improvement in eastern and southern African maize germplasm.

This study identifies two types of transition and transversion mutations. Transition mutations are nucleotide substitutions where a purine is replaced with another purine (A ↔ G) or a pyrimidine with another pyrimidine (C ↔ T), accounting for 77,700 occurrences. While transversion mutations are substitutions where a purine is replaced with a pyrimidine or vice versa (A or G ↔ C or T), occurred 39,087 times (Table 2).

**Table 2.**
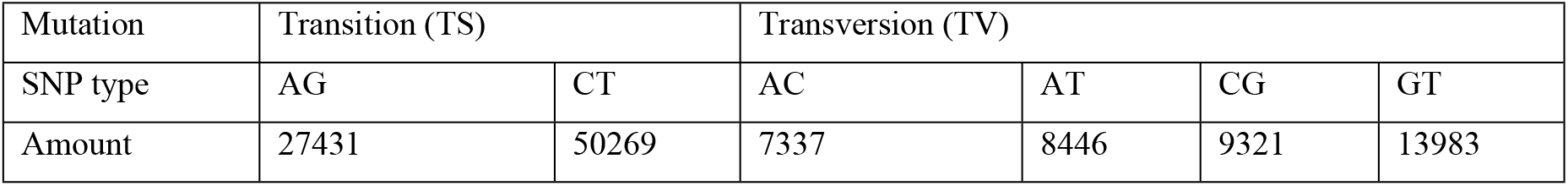
Single nucleotide polymorphism Mutations of Transition and Transversion types.

Pairwise genetic dissimilarity within the 2,555-maize doubled haploid lines ranged from 0 to 0.96, with average value of 0.91. The high genetic relatedness ranged from 0 to 0.2 and 74% of doubled haploid lines showed high genetic dissimilarity, ranged from 0.901 to 0.961. Additionally, pairwise kinship coefficients ranged from 0 to 1. Among the 2555 maize inbred lines, 0.3% of them were with kinship values close to zero, while 5.05% ranged between 0.501 to 0.60 and 61.8% fell between 0.801 to 0.90. The remaining 0.94% of maize doubled haploid lines had relative kinship ranged from 0.901 to 1(Figure 3).

**Figure 3.**
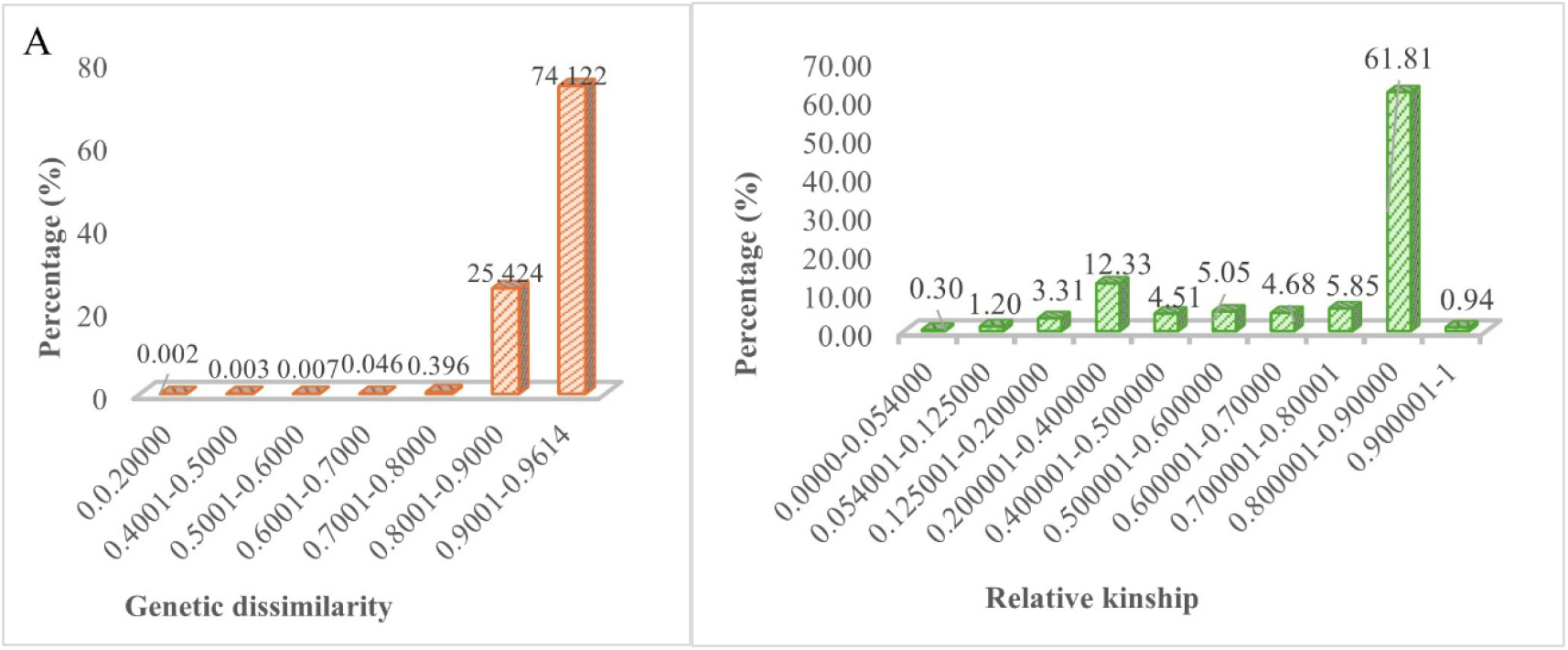
Dissemination of genetic dissimilarity and relative kinship among the 2,555 maize DH lines based on 2257 SNP

The population structure analysis using the sNMF (Sparse Non-negative Matrix Factorization) approach revealed an optimal number of genetic clusters (K) between 2 and 3, based on the minimum cross-entropy criterion (Figure 4). Visualization of individual ancestry proportions for K = 2 to 5 demonstrated clear sub-structuring within the DH panel (Figure 5). At K = 2, two distinct subpopulations emerged that corresponded closely to the known heterotic groups, with group A and group B represented by blue and red segments, respectively (Figure 6). This genetic differentiation aligned well with the breeding history of the DH lines.

**Figure 4.**
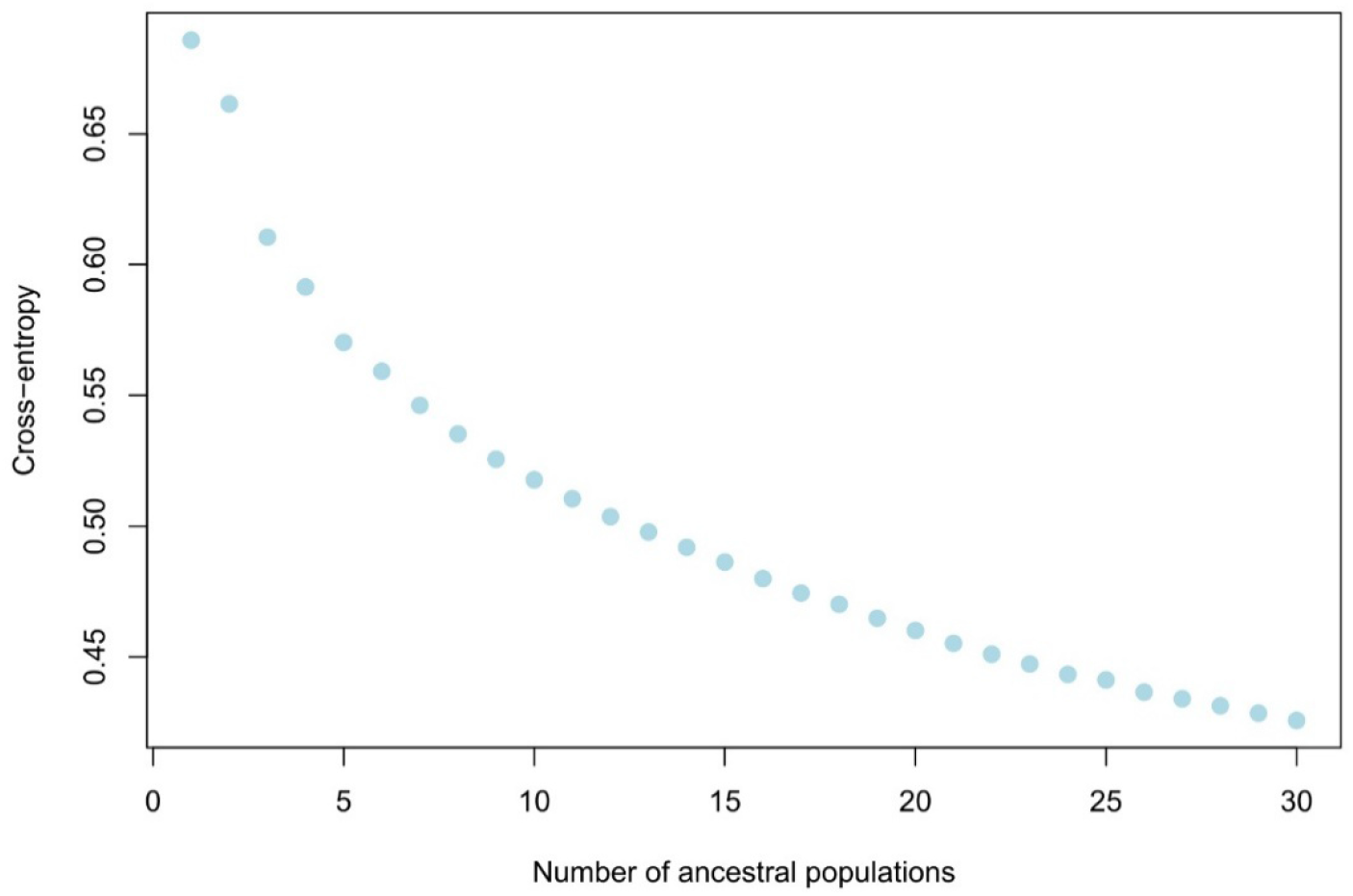
Mean values of the cross entropy estimated with the SNP dataset and computed across the 10 runs of sNMF performed for each K from K=2 to K=30.

**Figure 5.**
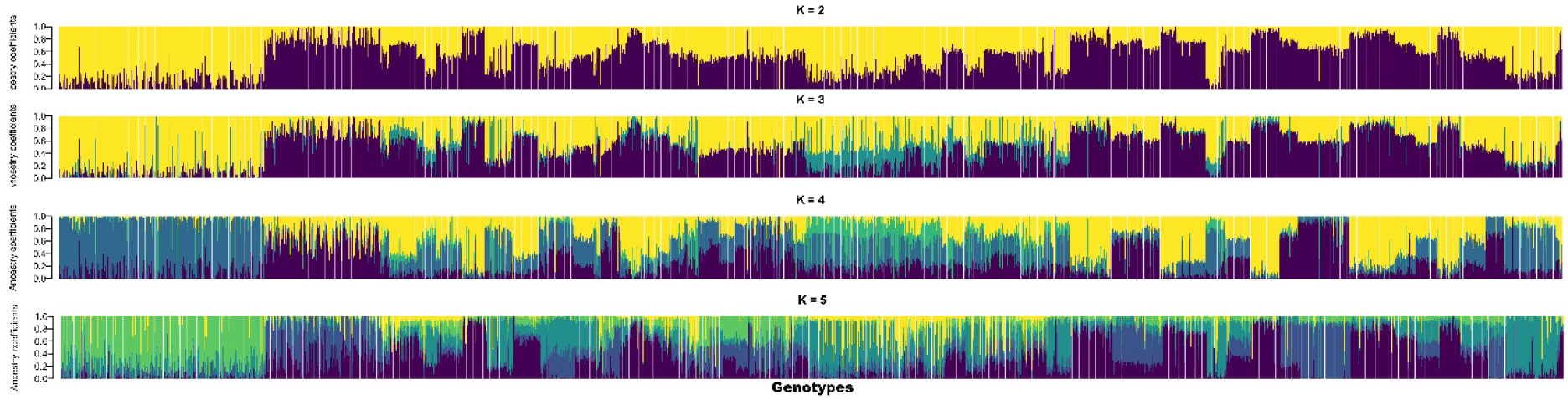
Population structure (from K = 2 to K = 5) of 2255 DH lines estimated with sNMF using the 2257 SNPs filtered for locus missingness and MAF at 0.05. Each individual is represented by a vertical bar, divided into K segments representing the proportion of genetic ancestry from the K clusters.

**Figure 6.**
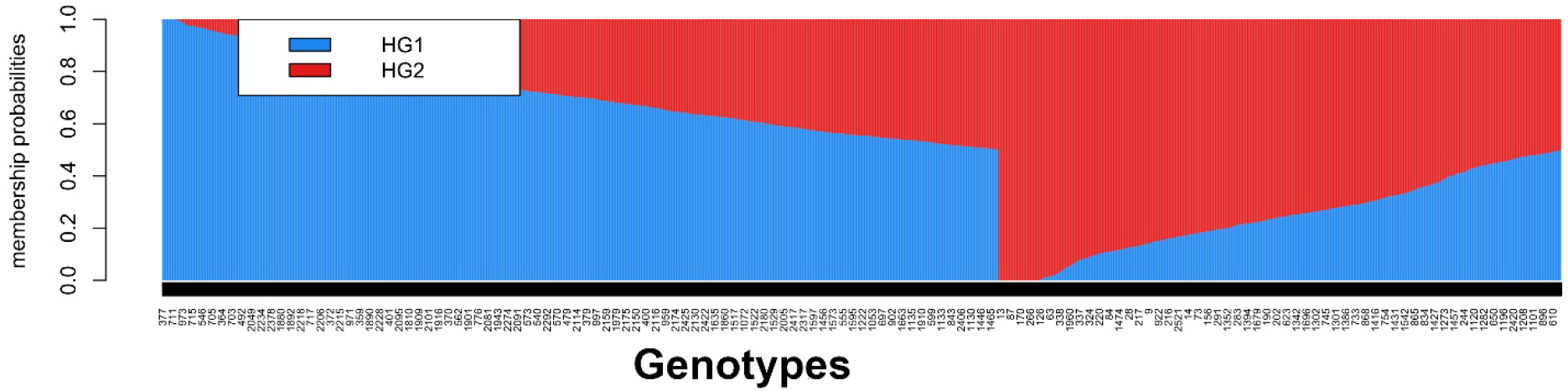
Analysis of population structure of 2555 maize doubled haploid lines within 2257 SNP With grouping based on heterotic groups when *K* = 2. The DH lines with blue, and red color representing Heterotic group subpopulations 1 (A) and 2 (B), respectively

In addition to population structure, the Principal Coordinate Analysis (PCoA) and hierarchical clustering method was also used to explore population structure. The clustering analysis resulted in two to three major distinct clusters, each containing a different number of inbred lines (Figure 7). Each sub-cluster was colored centered on the results of population structure. This classification reflects the genetic differentiation among the DH lines, with the majority of lines grouped into the larger clusters and fewer lines in the smaller clusters (Figure 7).

**Figure 7.**
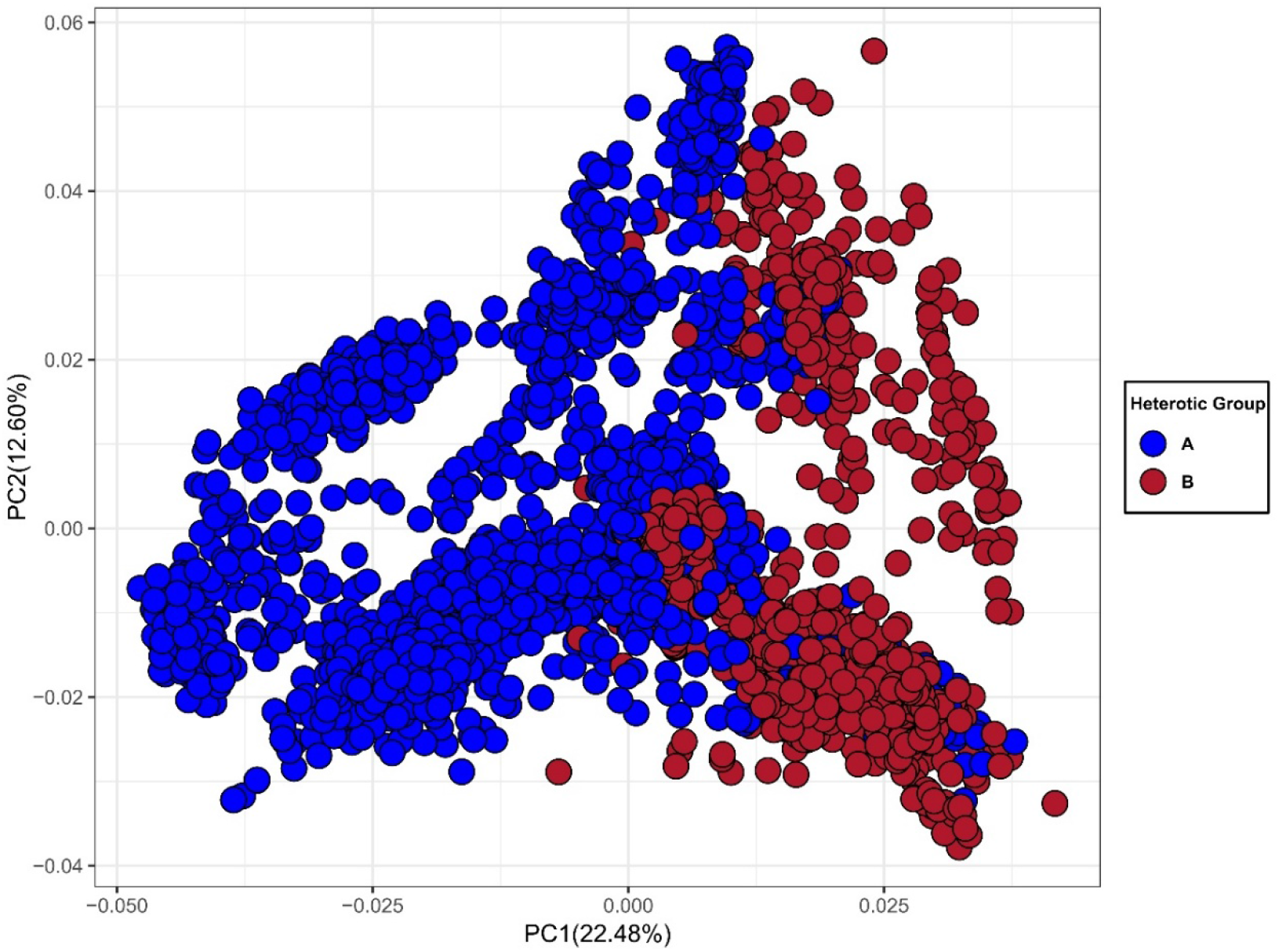
Principal coordinate analysis of 2555 maize DH lines with 2257 SNP markers. Color codes indicate the membership of the maize DH lines with heterotic group A (blue) and B (red).

The Principal Coordinate Analysis (PCoA) further supported the population structure results (Figure 8). In the maize doubled haploid lines, PC1 provides 22.48% of the total SNP variations, whereas PC2 accounts for 12.6% of the variation (Figure 7). DH lines grouped distinctly according to heterotic group assignments, with partial overlap between clusters, indicating shared ancestry among some genotypes. Similar grouping was also observed with hierarchical clustering for heterotic group A and B (Figure 8). The clear separation between the two heterotic groups highlights the genetic divergence that has been maintained through selective breeding.

**Figure 8.**
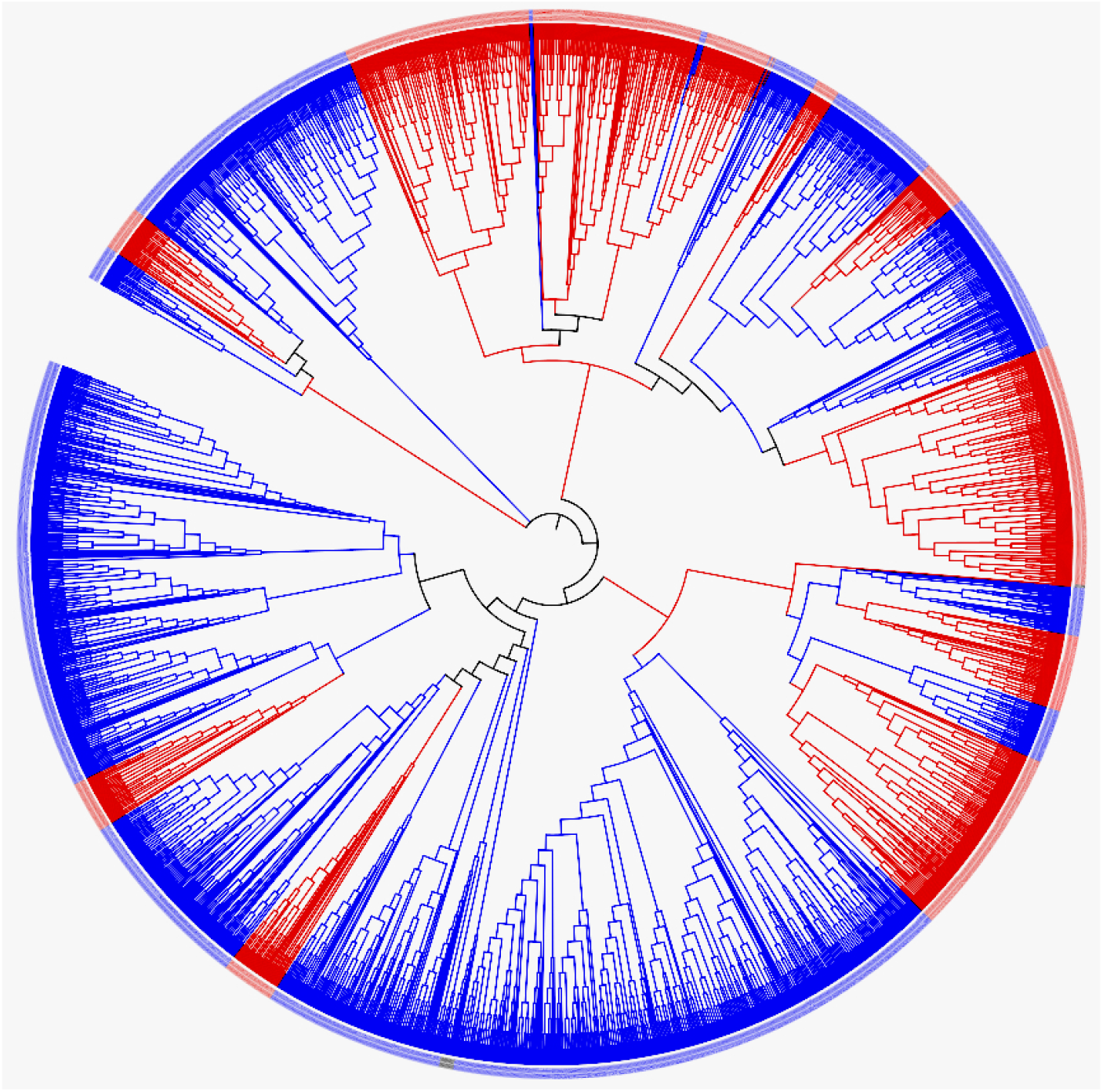
Hierarchical cluster of the 2555 DH lines, grouping is based on heterotic groups: The cluster with red color belongs to heterotic group B and blue color represents heterotic group A

### Analysis of Molecular Variance (AMOVA)

The AMOVA results (Table 3) partitioned the total molecular variance into components among and within populations. Differences among populations accounted for 36% of the total genetic variation, while 58% was attributed to differences among individuals within populations, and 6% was explained by variation within individuals. All variance components were highly significant (P = 0.001, based on 999 permutations), confirming the presence of strong genetic differentiation and extensive individual-level variation within the DH panel.

**Table 3:** Statistical Analysis of Molecular Variance (AMOVA)

| Source | DF | SS | MS | Est. Var. | %TV | P-value |
| --- | --- | --- | --- | --- | --- | --- |
| Among pops. | 86 | 101052.812 | 1175.033 | 18.992 | 36% | 0.001 |
| Among indiv. | 2468 | 160114.770 | 64.876 | 30.834 | 58% | 0.001 |
| Within indiv. | 2555 | 8195.500 | 3.208 | 3.208 | 6% | 0.001 |
| Total | 5109 | 29363.082 |  | 53.034 | 100% |  |

DF: degree of freedom, SS: sum of squares, MS: mean squares, Est. Var. estimate of variance, %TV percentage of the total variation and P-value are based on 999 permutations

Collectively, these diversity analyses demonstrate that the maize DH panel harbors moderate to high genetic diversity and exhibits distinct population structure consistent with heterotic group differentiation. The substantial within-population variability and low relatedness among most lines indicate that the panel provides a robust genetic resource for genome-wide association studies, genomic selection, and the identification of alleles associated with key agronomic and stress-resilience traits.

## Discussion

Genetic diversity serves as the fundamental resource for sustaining long-term genetic gain in tropical maize breeding. In sub-Saharan Africa, effective utilization of this diversity is critical for developing high-performing and climate-resilient hybrids adapted to heterogeneous production environments. The comprehensive analysis of 2,555 maize doubled haploid (DH) lines using 2,257 high-quality SNP markers revealed substantial genetic diversity and clear population structure consistent with the known heterotic grouping used in CIMMYT’s tropical maize breeding program. The mean gene diversity (0.38) and polymorphic information content (0.30) observed in this study indicate a moderate level of polymorphism, comparable to previous reports on tropical and subtropical maize germplasm [39]. The broad distribution of minor allele frequencies suggests that both common and rare alleles are well represented, ensuring the utility of this panel for downstream genomic analyses such as genome-wide association studies (GWAS) and genomic prediction [2,8].

The low observed heterozygosity (mean Ho = 0.04) and high fixation index (F = 0.89) confirmed the expected homozygosity typical of doubled haploid lines. This high genetic fixation provides an advantage for allele discovery and trait dissection since each genotype represents a stable and reproducible genetic entity. Despite this, the relatively wide range of gene diversity and PIC values indicates that a substantial proportion of allelic variation has been retained through the selection of genetically diverse parental sources during DH development [26].

To determine the types of genetic alterations taking place in these maize doubled haploid lines, transition (TS) and transversion (TV) mutations were examined. Transition mutations (A↔G and C↔T) were more frequent (77,700 occurrences), whereas transversion mutations (AC, AT, CG and GT) occurred 39,087 times, Purine-to-purine or pyrimidine-to-pyrimidine substitutions in genome are naturally stable, which may explain the higher frequency of transition mutations. In contrast, transversions, which involve more drastic base pair changes, are less common but may have more noticeable effects on fitness and phenotype. TS was higher than TV in this study. These findings are comparable to those previously reported by [15].

The relatedness between pairs of inbred lines was determined by the relative kinship coefficients, which varied 0 to 1. Low genetic diversity indicated the value of relatedness close to one, while high genetic diversity was indicated the value close to zero. The kinship percentage ranges from 0.801 to 0.90, accounting for 61.8% of all maize doubled haploid lines. This significant proportion indicates that these lines are more related to each other. Kinship results for the remaining 0.94% ranged from 0.901 to 1, suggesting doubled haploid lines that were extremely closely linked. With the majority of the lines showing moderate to high relatedness and a few genetically different lines that could be helpful for increasing the genetic base of breeding programs, these kinship coefficients emphasize the complicated genetic associations among the DH lines. However, this study indicates 5% proportion of kinship values were near to zero this result is less than the previous studies by [1,10,13,45], but greater than reported by [9,44].

Population structure analysis using sNMF and PCoA consistently identified two major genetic clusters corresponding to the established heterotic groups (A and B; Figure 6 and 7). These results are consistent with earlier reports showing that CIMMYT’s tropical maize germplasm maintains distinct but complementary heterotic patterns essential for maximizing heterosis and hybrid performance [2, 39]. The strong alignment between molecular clustering and heterotic group assignments underscores the efficiency of the breeding strategy in maintaining inter-group divergence while broadening genetic diversity within groups to support long-term genetic gain.

The analysis of molecular variance further confirmed significant population differentiation, with 36% of the total variance explained by differences among populations and 58% among individuals within populations. This pattern highlights the existence of both distinct heterotic pools and substantial within-group diversity, a feature that provides a broad genetic base for improvement of complex quantitative traits such as drought tolerance, yield stability, and resistance to major pests and diseases [23, 32]. The large proportion of intra-population variation emphasizes the potential of the DH panel for allele mining and recombination to enhance adaptive traits under diverse agroecological conditions.

Overall, the diversity patterns observed in this study confirm that the DH panel represents a robust and diverse genetic resource for tropical maize improvement. The presence of distinct population structure, high homozygosity, and substantial allelic richness makes the panel ideally suited for GWAS and genomic selection, where balanced diversity and structure are essential for accurate marker–trait association and prediction. Moreover, the results emphasize the effectiveness of CIMMYT’s long-term breeding strategy in maintaining heterotic group integrity while diversifying its germplasm base to support sustainable hybrid development for sub-Saharan Africa and other tropical regions [2, 22].

### Implications for breeding

The observed genetic diversity and structured differentiation among the 2,555 maize DH lines have important implications for tropical maize breeding in sub-Saharan Africa. The presence of two distinct heterotic groups, coupled with considerable within-group variability, provides a solid foundation for designing high-yielding and stress-resilient hybrids. The broad allelic variation captured across loci enhances opportunities to identify favorable alleles for drought tolerance, insect-pest resistance, and grain quality traits [19, 28]. The moderate differentiation and extensive within-population diversity indicate that sufficient genetic variability remains available for recombination and long-term selection gain. These results also validate CIMMYT’s breeding strategy of maintaining heterotic group integrity while diversifying the genetic base to adapt to evolving climatic and biotic challenges. The DH panel, therefore, represents a valuable resource for genomic selection, GWAS, and integrative omics research aimed at accelerating genetic improvement and ensuring food security in tropical maize systems of SSA.

## Conclusion

The observed genetic diversity and population structure among the 2,555 maize DH lines provide a strong genomic foundation for trait dissection and molecular breeding. The well-defined heterotic groups, combined with substantial within-group variability, offer unique opportunities to identify novel alleles and genomic regions associated with key adaptive and productivity traits. Building on this genomic framework, integrating transcriptomic and metabolomic data will enable a deeper understanding of gene networks and metabolic pathways underpinning native resistance to fall armyworm, drought tolerance, and other stress-related traits, ultimately accelerating genetic gain in tropical maize breeding programs.

## Author Contributions

Conceptualization, funding acquisition, project & resources administration

## Acknowledgments

The authors are grateful to the International Maize and Wheat Improvement Center (CIMMYT) scientists and technicians who generated the germplasm, and highly appreciate the technical support received from the staff members affiliated to CIMMYT maize research station in Kiboko, Kenya, Addis Ababa Science and Technology University and Institution support seed system in Ethiopia.

## Supporting Information

**Table 1:** List of maize single-cross source populations, pedigree, heterotic group, and number of DH lines used in the study

## References

1. Adu, G. B., Awuku, F. J., Garcia-Oliveira, A. L., Amegbor, I. K., Nelimor, C., Nboyine, J., Karikari, B., Atosona, B., Manigben, K. A., & Aboydana P. A. (2024). DArTseq-based SNP markers reveal high genetic diversity among early generation fall armyworm tolerant maize inbred lines. PLoS ONE 19(4): e0294863. 10.1371/journal.pone.0294863

2. Beyene, Y., Gowda, M., Olsen, M., Robbins, K. R., Pérez-Rodríguez, P., Alvarado, G., Dreher, K., Gao, S. Y., Mugo, S., Prasanna, B. M., & Crossa, J. (2019). Empirical comparison of tropical maize hybrids selected through genomic and phenotypic selections. Frontiers in Plant Science, 10, 1502. 10.3389/fpls.2019.01502

3. Beyene, Y., Gowda, M., Pérez-Rodríguez, P., Olsen, M., Robbins, K. R., Burgueño, J., Rincon, F., Monson, M., Nair, S. K., Babu, R., Das, B., Prasanna, B. M & Crossa, J. (2021). Application of genomic selection at the early stage of breeding pipeline in tropical maize. Frontiers in Plant Science, 12, 685488. 10.3389/fpls.2021.685488

4. Cairns, J. E., Crossa, J., Zaidi, P. H., Grudloyma, P., Sanchez, C., Araus, J. L., Thaitad, S., Makumbi, D., Magorokosho, C., Bänziger, M., Menkir, A., Hearne, S., & Atlin, G. N. (2013). Identification of drought, heat, and combined drought and heat tolerant donors in maize. Crop Science, 53(4), 1335–1346. 10.2135/cropsci2012.09.0545

5. Chaikam, V., Martinez, L., Melchinger, A. E., Schipprack, W., & Boddupalli, P. M. (2016). Development and validation of red root marker-based haploid inducers in maize. Crop Science, 56(4), 1678–1688. 10.2135/cropsci2015.09.0561

6. Chang, C. C., Chow, C. C., Tellier, L. C., Vattikuti, S., Purcell, S. M., & Lee, J. J. (2015). Second-generation PLINK: rising to the challenge of larger and richer datasets. Gigascience, 4(1), s13742–015. 10.1186/s13742-015-0047-8

7. Chao, A., & Shen, T. J. (2003). Nonparametric estimation of Shannon’s index of diversity when there are unseen species in sample. Environmental and ecological statistics, 10(4), 429–443. 10.1023/A:1026096204727

8. Crossa, J., Pérez-Rodríguez, P., Cuevas, J., Montesinos-López, O., Jarquín, D., De Los Campos, G., Burgueño, J., González-Camacho, J. M., Pérez-Elizalde, S., Beyene, Y., Dreisigacker, S., Singh, R., Zhang, X., Gowda, M., Roorkiwal, M., Rutkoski, J., & Varshney, R. K. (2017). Genomic selection in plant breeding: methods, models, and perspectives. Trends in plant science, 22(11), 961–975. 10.1016/j.tplants.2017.08.011

9. Dao, A., Sanou, J., Mitchell, S.E., Gracen, V., & Danquah, E. Y. (2014). Genetic diversity among INERA maize inbred lines with single nucleotide polymorphism (SNP) markers and their relationship with CIMMYT, IITA, and temperate lines. BMC Genetics, 15, 1–14. 10.1186/s12863-014-0127-2

10. de Faria, S.V., Zuffo, L.T., Rezende, W. M., Caixeta, D. G., Pereira, H. D., Azevedo, C. F., & Delima R. O. (2022). Phenotypic and molecular characterization of a set of tropical maize inbred lines from a public breeding program in Brazil. BMC Genomics, 23, 1–17. 10.1186/s12864-021-08127-7

11. Elshire, R. J., Glaubitz, J. C., Sun, Q., Poland, J. A., Kawamoto, K., Buckler, E. S., & Mitchell, S. E. (2011). A robust, simple genotyping-by-sequencing (GBS) approach for high diversity species. PloS one, 6(5), e19379. 10.1371/journal.pone.0019379

12. Erenstein, O., Jaleta, M., Sonder, K., Mottaleb, K., & Prasanna, B. M. (2022). Global maize production, consumption and trade: Trends and R&D implications. Agriculture & Food Security, 11, Article 25. 10.1186/s40066-022-00374-5

13. Ertiro, B.T., Semagn, K., Das, B., Olsen, M., Labuschagne, M., Worku, M., Wegay D., Azmach G., Ogugo V., Keno T., Abebe B., & Chibs T. (2017). Genetic variation and population structure of maize inbred lines adapted to the mid-altitude sub-humid maize agro-ecology of Ethiopia using SNP markers. BMC Genomics, 18, 1–11. 10.1186/S12864-017-4173-9

14. Food and Agriculture Organization of the United Nations (2024). Crop Prospects and Food Situation – Tri-annual Global Report No. 1, March 2024. Rome: FAO

15. Fufa, T.W., Abtew, W. G., Amadi, C. O., & Oselebe, H. O. (2022). DArTSeq SNP-based genetic diversity and population structure studies among taro [(Colocasia esculenta L.) Schott] accessions sourced from Nigeria and Vanuatu. PLOS ONE, 17(11): e0269302. 10.1371/journal.pone.0269302

16. Gaytán-Pinzón, G. P., Sandoval-Castro, E., Peinado-Fuentes, L. A., Valenzuela-Apodaca, J. P., Cruz-Mendívil, A., & Calderón-Vázquez, C. L. (2022). Genetic diversity of subtropical double-haploid maize lines selected for high oil content. Agronomy Journal, 114(5), 2715–2727. 10.1002/agj2.20909

17. Hallauer, A. R., Carena, M. J., & Miranda Filho, J. D. (2010). Quantitative genetics in maize breeding (Vol. 6). Springer Science & Business Media.

18. Jenkins, M. T., & Brunson, A. M. (1932). Methods of testing inbred lines of maize in crossbred combinations. Agronomy Journal, 24(7), 523–530. 10.2134/agronj1932.00021962002400070004x

19. Kamweru, I., Beyene, Y., Anani, B., Adetimirin, V. O., Prasanna, B. M., & Gowda, M. (2023). Hybrid breeding for fall armyworm resistance: Combining ability and hybrid prediction. Plant breeding= Zeitschrift fur Pflanzenzuchtung, 142(5), 607–620. 10.1111/pbr.13129

20. Kilian, A., Wenzl, P., Huttner, E., Carling, J., Xia, L., Blois, H., Caig, V., Heller-Uszynska, K., Jaccoud, D., Hopper, C., Aschenbrenner-Kilian, M., Evers, M., Peng, K., Cayla, C., Hok, P., & Uszynski, G. (2012). Diversity arrays technology: A generic genome profiling technology on open platforms. In T. Kantety & A. R. J. Mooney (Eds.), Data production and analysis in population genomics: Methods and protocols (Vol. 888, pp. 67–89). Totowa, NJ: Humana Press. 10.1007/978-1-61779-870-2_5

21. Lu, H., & Bernardo, R. (2001). Molecular marker diversity among current and historical maize inbreds. Theoretical and Applied Genetics, 103(4), 613–617. 10.1007/s001220100575

22. Mageto, E.K., Magorokosho, C., Suresh, L. M., Olsen, M. S., Crossa, J., Semagn, K., & Beyene, Y. (2022). Genetic diversity of CIMMYT maize inbred lines based on SNP markers. Genetic Resources and Crop Evolution, 69(6), 1457–1474. 10.1007/s10722-021-01309-8

23. Masuka, B., Atlin, G. N., Olsen, M., Magorokosho, C., Labuschagne, M., Crossa, J., Bänziger, M., Pixley, K. V., Vivek, B. S., Makumbi, D., Tarekegne, A., Das, B., Zaman-Allah, M. & Cairns, J. E. (2017). Gains in maize genetic improvement in Eastern and Southern Africa: I. CIMMYT hybrid breeding pipeline. Crop Science, 57(1), 168–179. 10.2135/cropsci2016.05.0343

24. Matsuoka, Y., Vigouroux, Y., Goodman, M. M., Sanchez G. J., Buckler, E., & Doebley, J. (2002). A single domestication for maize shown by multilocus microsatellite genotyping. Proceedings of the National Academy of Sciences of the United States of America, 99(9), 6080–6084. 10.1073/pnas.052125199

25. Melchinger, A. E., & Gumber, R. K. (1998). Overview of heterosis and heterotic groups in agronomic crops. Concepts and breeding of heterosis in crop plants, 25, 29–44. Crop Science Society of America. 10.2135/cssaspecpub25.c3

26. Menkir, A., Chikoye, D., Tofa, A. I., Fagge, A. A., Dahiru, R., Solomon, R., Ademulegun, T., Omoigui, L., Aliyu, K. T., & Kamai, N. (2020). Mitigating Striga hermonthica parasitism and damage in maize using soybean rotation, nitrogen application, and Striga-resistant varieties in the Nigerian savannas. Experimental agriculture, 56(4), 620–632. 10.1017/S0014479720000198

27. Mukiti, H. M., Badu-Apraku, B., Abe, A., Adejumobi, I. I., & Derera, J. (2025). Optimizing breeding strategies for early-maturing white maize through genetic diversity and population structure. PloS one, 20(2), e0316793. 10.1371/journal.pone.0316793

28. Noel, S. D. (2024). Developing a Research-Based Framework for Teaching Undergraduates with Archives. Simmons University.

29. Ogugo, V., Semagn, K., Beyene, Y., Runo, S., Olsen, M., & Warburton, M. L. (2015). Parental genome contribution in maize DH lines derived from six backcross populations using genotyping by sequencing. Euphytica, 202(1), 129–139. 10.1007/s10681-014-1290-5

30. Peakall, R. & Smouse, P. E. (2012). GenAlEx 6.5: Genetic analysis in Excel. Population genetic software for teaching and research an update. Bioinformatics 28, 2537–2539

31. Peakall, R., & Smouse, P. E. (2006). GENALEX 6: Genetic analysis in Excel. Population genetic software for teaching and research. Molecular Ecology Notes, 6(1), 288–295. 10.1111/j.1471-8286.2005.01155.x

32. Prasanna, B. M., Burgueño, J., Beyene, Y., Makumbi, D., Asea, G., Woyengo, V., Olsen, M., Das, B., Worku, M., Shikuku, K., Kasozi, S., Mugo, S., Tarekegne, A., Gowda, M., Bright, J. M., Crossa, J., & Cairns, J. E. (2022). Genetic trends in CIMMYT’s tropical maize breeding pipelines. Scientific Reports, 12(1), 20110. 10.1038/s41598-022-24561-2

33. Prasanna, B. M., Cairns, J. E., Zaidi, P. H., Beyene, Y., Makumbi, D., Gowda, M., Magorokosho, C., Zaman-Allah, M., Olsen, M., Das, A., Worku, M., Gethi, J., Vivek, B. S., Nair, S. K., Rashid, Z., Vinayan, M. T., Issa, A. B., San Vicente, F., Dhliwayo, T., & Zhang, X. (2021). Beat the stress: breeding for climate resilience in maize for the tropical rainfed environments. Theoretical and Applied Genetics, 134(6), 1729–1752. 10.1007/s00122-021-03773-7

34. Prasanna, B. M., Chaikam, V., & Mahuku, G. (2012). Doubled haploid technology in maize breeding: theory and practice. CIMMYT.

35. Prigge, V., Xu, X., Li, L., Babu, R., Chen, S., Atlin, G. N., & Melchinger, A. E. (2012). New insights into the genetics of in vivo induction of maternal haploids, the backbone of doubled haploid technology in maize. Genetics, 190(2), 781–793. 10.1534/genetics.111.133066

36. Pritchard, J. K., Stephens, M., & Donnelly, P. (2000). Inference of population structure using multilocus genotype data. Genetics, 155(2), 945–959. 10.1093/genetics/155.2.945

37. R Core Team. (2023). R: A language and environment for statistical computing. R Foundation for Statistical Computing. https://www.R-project.org/

38. Rawlings, J. O., & Thompson, D. L. (1962). Performance level as criterion for the choice of maize testers. Crop Science, 2(3), 217–220. 10.2135/cropsci1962.0011183X000200030012x

39. Semagn, K., Magorokosho, C., Vivek, B. S., Makumbi, D., Beyene, Y., Mugo, S., Prasanna, B. M., & Warburton, M. L. (2012). Molecular characterization of diverse CIMMYT maize inbred lines from eastern and southern Africa using single nucleotide polymorphic markers. BMC Genomics, 13(1), 113. 10.1186/1471-2164-13-113

40. Senior, M. L., Chin, E. C. L., Lee, M., Smith, J. S. C., & Stuber, C. W. (1996). Simple sequence repeat markers developed from maize sequences found in the GENBANK database: map construction. Crop Science, 36(6), 1676–1683.

41. Shiferaw, B., Prasanna, B. M., Hellin, J., & Bänziger, M. (2011). Crops that feed the world 6. Past successes and future challenges to the role played by maize in global food security. Food security, 3(3), 307–327. 10.1007/s12571-011-0140-5

42. Smith, J. S. C., Chin, E. C. L., Shu, H., Smith, O. S., Wall, S. J., Senior, M. L., Mitchell, S. E., Kresovich, S., & Ziegle, J. (1997). An evaluation of the utility of SSR loci as molecular markers in maize (Zea mays L.): comparisons with data from RFLPs and pedigree. Theoretical and Applied Genetics, 95(1), 163–173. 10.1007/s001220050544

43. Warburton, M. L., Reif, J. C., Frisch, M., Bohn, M., Bedoya, C., Xia, X. C., Crossa, J., Franco, J., Hoisington, D., Pixley, K., Taba, S., & Melchinger, A. E. (2008). Genetic diversity in CIMMYT nontemperate maize germplasm: Landraces, open pollinated varieties, and inbred lines. Crop Science, 48(2), 617–624. 10.2135/cropsci2007.09.0511

44. Wu, X., Li, Y., Shi, Y., & Song, Y. (2014). Fine genetic characterization of elite maize germplasm using high-throughput SNP genotyping. Theoretical and Applied Genetics, 128(3), 621–631. 10.1007/s00122-013-2246-y

45. Wu, Y., San Vicente, F., Huang, K., Dhliwayo, T., Costich, D. E., Semagn, K., Sudha, N., Olsen, M., Prasanna, B. M., Zhang, X. & Babu, R. (2016). Molecular characterization of CIMMYT maize inbred lines with genotyping-by-sequencing SNPs. Theoretical and Applied Genetics, 129(4), 753–765.

46. Xia, H. J., & Yang, G. (2005). Inositol 1, 4, 5-trisphosphate 3-kinases: functions and regulations. Cell research, 15(2), 83–91. 10.1038/sj.cr.7290275

47. Yan, J., Shah, T., Warburton, M. L., Buckler, E. S., McMullen, M. D., & Crouch, J. (2009). Genetic characterization and linkage disequilibrium estimation of a global maize collection using SNP markers. PloS one, 4(12), e8451. 10.1371/journal.pone.0008451

